# SynergyFinder Plus: Toward Better Interpretation and Annotation of Drug Combination Screening Datasets

**DOI:** 10.1101/2021.06.01.446564

**Authors:** Shuyu Zheng, Wenyu Wang, Jehad Aldahdooh, Alina Malyutina, Tolou Shadbahr, Ziaurrehman Tanoli, Alberto Pessia, Jing Tang

## Abstract

Combinatorial therapies have been recently proposed to improve the efficacy of anticancer treatment. The SynergyFinder R package is a software used to analyze pre-clinical drug combination datasets. Here, we report the major updates to the SynergyFinder R package for improved interpretation and annotation of drug combination screening results. Unlike the existing implementations, the updated SynergyFinder R package includes five main innovations. (1) We extend the mathematical models to higher-order drug combination data analysis and implement dimension reduction techniques for visualizing the synergy landscape. (2) We provide a statistical analysis of drug combination synergy and sensitivity with confidence intervals and *P* values. (3) We incorporate a synergy barometer to harmonize multiple synergy scoring methods to provide a consensus metric for synergy. (4) We evaluate drug combination synergy and sensitivity to provide an unbiased interpretation of the clinical potential. (5) We enable fast annotation of drugs and cell lines, including their chemical and target information. These annotations will improve the interpretation of the mechanisms of action of drug combinations. To facilitate the use of the R package within the drug discovery community, we also provide a web server at www.synergyfinderplus.org as a user-friendly interface to enable a more flexible and versatile analysis of drug combination data.

## Introduction

Many complex diseases including cancer and infectious diseases develop drug resistance [1,2]. To achieve more sustainable clinical efficacy, multi-targeted drug combinations have been proposed to tackle disease signaling pathways more systematically [3]. With the advances in high-throughput drug screening technologies, an increasing number of drug combinations are being tested in multiple disease models, such as cancer cell lines and patient-derived primary cultures. These preclinical drug combination experiments are conducted to identify the most synergistic and effective drug combinations that result in improved cellular responses, such as cell viability and toxicity. With the potential hits prioritized from screening, more functional studies are warranted to elucidate the mechanisms of action of the drug interactions, leading to the identification of drug combination response biomarkers that are critical for stratifying patients for more targeted therapies.

For the evaluation of the potential of a drug combination, mathematical and statistical models are needed to characterize the expectation if the drugs are not interactive; thereafter the boosting effects of the drug combination can be formally quantified. There are currently four major synergy models: highest single agent (HSA) [4], Loewe additivity (LOEWE) [5], Bliss independence (BLISS) [6], and zero interaction potency (ZIP) [7]. However, the overall effects of a drug combination may not lead to sufficient efficacy, despite a strong degree of interaction. Therefore, it is recommended to evaluate both synergy (*i.e*., the degree of interactions) and sensitivity (*i.e*., the overall treatment efficacy) simultaneously [8]. Previously developed tools that can analyze drug combination synergy include COMPUSYSN [9], BRAID [10], Combenfit [11], SynergyFinder [12], and Synergy [13]. However, these tools are designed mainly for analyzing two-drug combinations only. More recently, the SynergyFinder2 tool [14] has been developed, which extends to three-drug combinations. However, only the HSA model was appropriately constructed for its implementation. The key differences in the modeling are summarized in Table S1. Furthermore, the visualization of high-order drug combinations becomes non-trivial, as none of the existing visualization methods can deal with combinations of three or more drugs [15]. More importantly, there is a lack of implementation tools that can harmonize multiple synergy models to derive a consensus about the degree of interactions [16].

To address these limitations, we provide a major update of the SynergyFinder R package that enables novel functions, including (1) formal mathematical models to assess the synergy of high-order drug combinations and visualization of the degree of synergy using a dimension reduction technique; (2) formal statistical methods to evaluate the significance of synergy; (3) implementation of the synergy barometer to systematically compare the results from different synergy models; and (4) implementation of a synergy-sensitivity (SS) plot to evaluate the potential of a drug combination unbiasedly. (5) Finally, we develop data annotation tools to retrieve the pharmacological and molecular profiles of the drugs and cells, facilitating the discovery of mechanisms of action of the most synergistic and effective drug combinations. We provide a new website at www.synergyfinderplus.org to allow users to upload their experimental results and run all the analyses with the R package in the backend. All functions and source codes are freely accessible to academic users (**Figure 1**).

**Figure 1.**
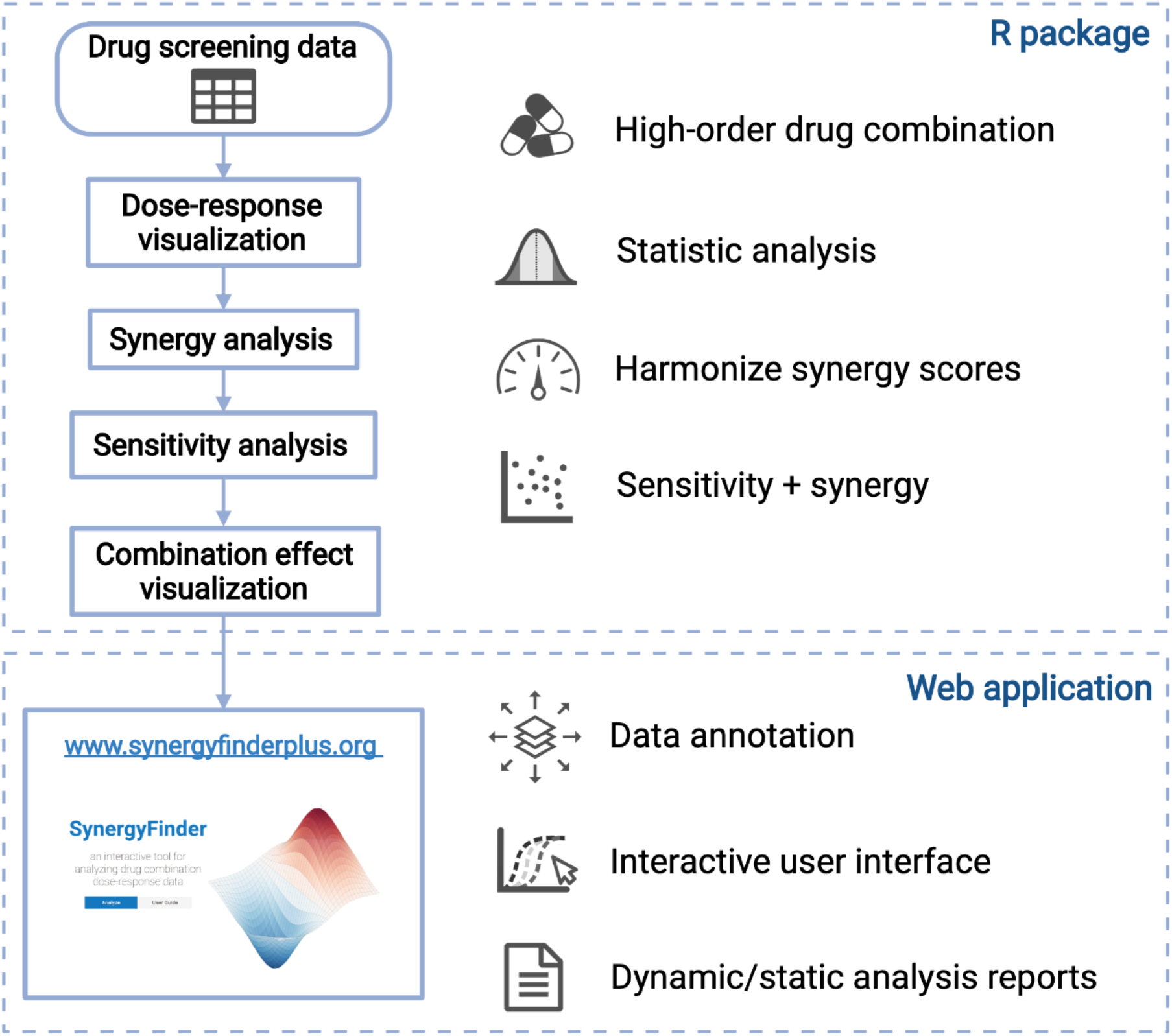
Summary of the SynergyFinder Plus workflow. We highlight the new features including: (1) the ability to analyze drug combinations of more than two drugs, (2) tailored statistical testing to account for uncertainty for data with replicates, (3) harmonized visualization of multiple synergy score metrics, (4) visualization of the relationship between sensitivity and synergy scores, and (5) a data annotation web server to query cell line and drug information.

## Method

### Synergy models for high-order drug combinations

Considering the response of a drug to be measured as a % inhibition *y* that ranges from 0 to 1, a higher value indicates a better efficacy. For a combination that involves *n* drugs, the observed combination response is denoted as *y_c_*, whereas the observed monotherapy response of its constituent drugs is *y*_*i,i*=1,…*n*_. The expected combination response is determined by the assumption of non-interaction. Currently there are four major synergy reference models [16,17]. For the HSA model, the expected response is the highest monotherapy response, that is, *y_HSA_* = max(*y*_1_,…, *y_i_*,…, *y_n_*). For the BLISS model, the expected response is derived from the probabilistic independence of the monotherapy effects, that is, *y_BLISS_* = 1 − Π_*i*_(1 − *y_i_*). For the LOEWE model, the expected response satisfies 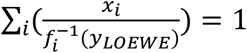, where *x_i_* is the dose of each constituent drug in the combination, and 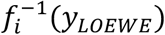 is the inverse function of the dose-response curve. For the ZIP model, the expected response satisfies 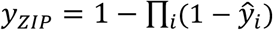, where 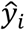 is the predicted dose response of the monotherapy by a monotonically increasing curve fitting model 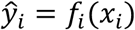. For example, a common choice for modeling drug dose-response curves is the four-parameter log-logistic model 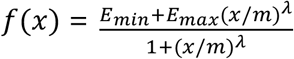, where *E_min_*, *E_max_*, *m*, and *λ* are the minimal and maximal responses, IC50, and slope of the dose-response curve, respectively.

Accordingly, the multi-drug synergy score for the observed combination response *y_c_* can be determined as follows:

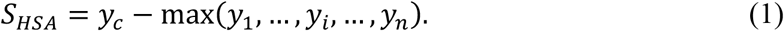

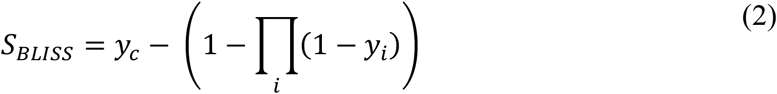

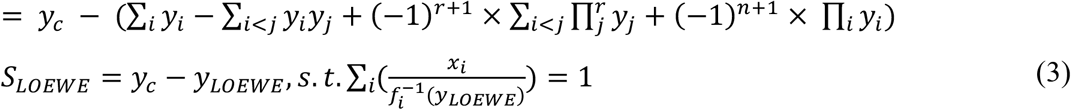

To determine the ZIP-based synergy score, *y_c_* needs to be replaced with the predicted average response 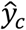 given by the curve fitting models 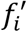 to make it comparable to *y_ZIP_*:

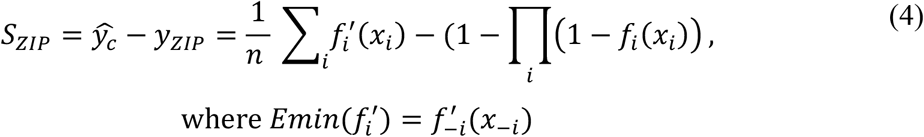

Namely, 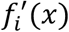 stands for the log-logistic model defined for the combination response at dose *x* of drug *i* when the other drugs are present. Furthermore, *Emin* of 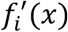 is determined by 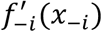, which is the fitted curve of the combination response while drug *i* is absent. Note that the ZIP model captures the shift of potency for a drug combination in comparison to its monotherapy drugs; therefore, the ZIP model compares the difference between fitted models for the drug combination 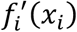 and for the monotherapy drug *f_i_*(*x_i_*).

### Visualization of higher-order synergy and sensitivity scores

The commonly utilized synergy or sensitivity landscape models are not directly applicable for the visualization of higher-order drug combinations. It is non-trivial to determine the coordinates of the multi-dimensional dose vectors in a two-dimensional space where similar dose vectors remain proximal to each other. In this study we propose a dimension reduction technique that is based on multidimensional scaling, similar to the recent application in transforming numerical data into images [18]. For a drug combination in an *n*-dimensional dose space, with its coordinates of *X* = (*x*_1_,… *x_n_*), we consider their rankings *R* = (*r*_1_,… *r_n_*). For example, a combination *X = (0.1, 10, 1000)* nM in three-dimensional dose space has a ranking vector R = (1, 3, 6), suggesting that 0.1 nM is the minimum dose in the tested dose range of drug 1, 10 nM is the third smallest dose of drug 2, and 1000 nM is the sixth dosage of drug 3. Euclidean distance is a common choice for quantifying the similarity between two dose ranking vectors. We utilize multi-dimensional scaling to minimize the error of the pairwise distance in a two-dimensional space in which the synergy and sensitivity scores can be visualized as a synergy landscape (**Figure 2**). Using the dose rankings as the input for multi-dimensional scaling can ensure that the resulting two-dimensional coordinates are equally distanced among the neighboring dose conditions, thus making the visualization easier to interpret. Furthermore, for the case of two-drug combinations, the algorithm can converge to the actual dose rankings, thus preserving the consistency across all orders of combinations. We also develop functions to visualize the synergy and sensitivity scores for a specific dose condition in a grid of bar plots, which can help identify the most synergistic and sensitive areas in the synergy landscape.

**Figure 2.**
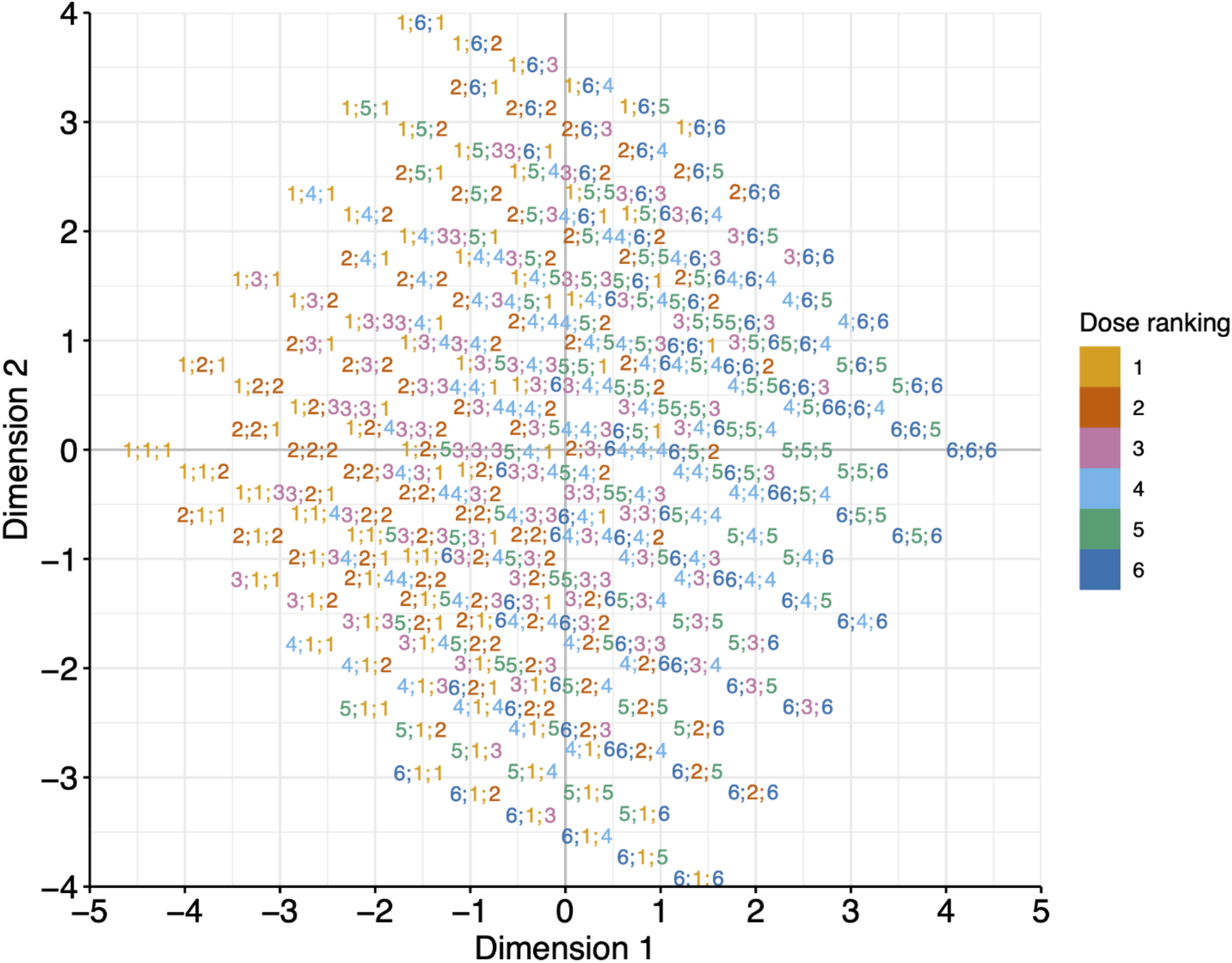
Dimension reduction for the visualization of high-order drug combinations. In this example, the two-dimensional coordinates of each three-drug dose combination are determined by multidimensional scaling based on the similarity of their dose ranges, after which the synergy scores and sensitivity scores can be visualized on the map. Each dot in the map is annotated with its associated dose rankings for the three drugs (*r*_1_; *r*_2_; *r*_3_). For example, a dot with a label 4;1;2 shows a combination involves 4^th^ dose of drug 1, 1^st^ dose of drug 2, and 2^nd^ dose of drug 3.

### Statistical analysis

The statistical significance of synergy can be defined at a single dose or at the whole dose matrix level. Consider a two-drug combination experiment as an example. We assume that the replicates of drug responses are measured independently within each dose in the dose matrix (**Figure 3**). At each dose level, we propose using a bootstrapping approach to determine the confidence intervals of the synergy scores. To determine a bootstrap dose-response matrix, we sample the responses for each dose combination from a normal distribution *N*(*μ, σ*), where *μ* and *σ* are the mean and standard deviation of the replicates. The synergy scores for the HSA and BLISS models can be derived simply by comparing the bootstrapped responses at the dose combination and at their corresponding monotherapy doses (*i.e*., using Equations (1) and (2)). However, for the LOEWE and ZIP models [7], bootstrapped synergy scores will be derived using the curve fitting over the whole dose matrix (*i.e*., Equations (3) and (4)).

**Figure 3.**
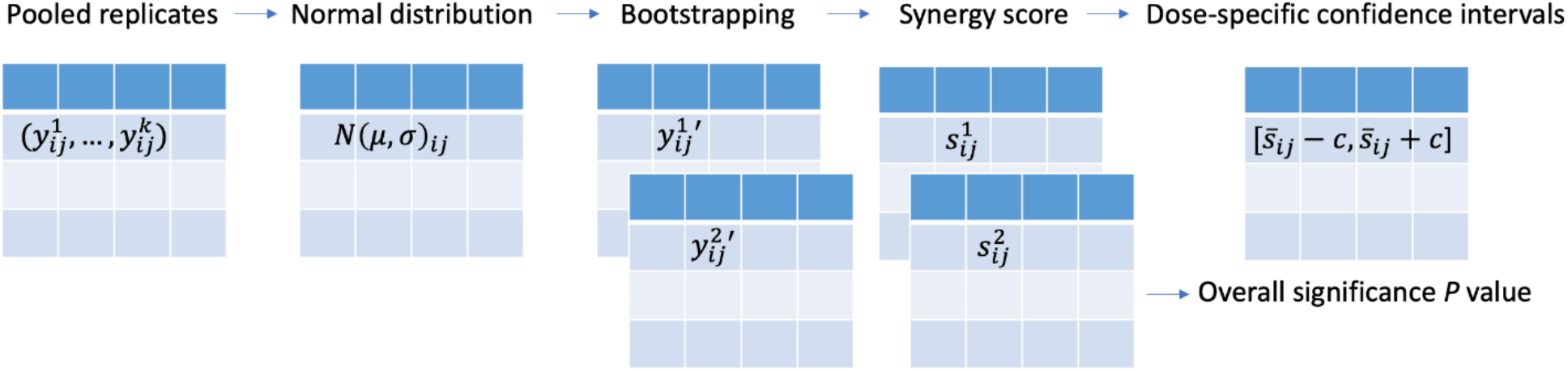
Statistical testing of synergy scores. Dose-specific confidence intervals and *P* values for the average synergy score can be derived using bootstrapping of the dose-response matrix.

Suppose that *B* bootstrap samples are drawn from the replicates and suppose that *S*_1_,…, *S_B_* are the estimates of the synergy scores determined from these bootstrap samples, with a mean of 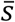. We determine the bootstrap standard error (SE) as follows:

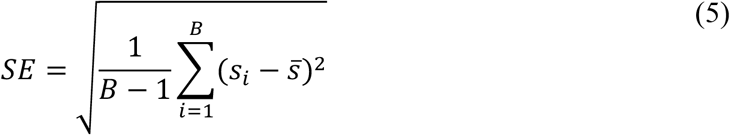

The 95% confidence interval for the synergy score is approximately

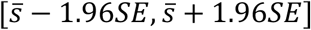

At the whole dose matrix level, we provide an empirical *P* value to assess the significance of the difference between the estimated average synergy score over the whole dose matrix and the expected synergy score of zero under the null hypothesis of non-interaction.

Letting 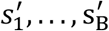 are the estimates of the average synergy scores over the whole dose matrix from the bootstrap samples, the *P* value is [19]:

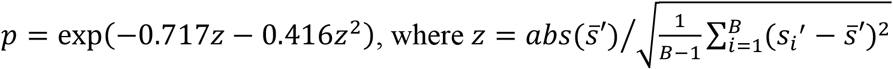

For a scenario in which no replicates are available, the confidence interval cannot be determined at the dose level. However, the *P* value of the average synergy score of the whole dose-response matrix can still be derived by pooling the synergy scores together and then comparing them to zero. For both scenarios, we report the overall *P* values of synergy and the confidence intervals at the dose-specific level when replicates are available. We test the accuracy of the bootstrapping with the simulation (File S1) and show that the *P* values for synergy scores determined from bootstrap are highly correlated with the true *P* values (Figure S1).

### Harmonization of multiple synergy models

As the SynergyFinder R package provides four synergy models (*i.e*., HSA, BLISS, LOEWE, and ZIP) that differ in their null hypotheses of non-interaction, the synergy scores do not necessarily coincide with each other for a given dataset. To provide a more harmonized representation of the synergy score results, we introduce a novel tool to enable a more systematic comparison of the synergy scores. We leverage the implementation of the four models to derive the expected response of non-interaction (*i.e*., *y_HSA_*, *V_BLISS_*, *Z_LOEWE_*, and *y_ZIP_*). Because all the expected responses share the same unit as the observed response (*i.e*., % inhibition), we may utilize a synergy barometer to compare these values. The expected and observed responses for a given combination at a specific dose are positioned on the same scale. With such a tool as the synergy barometer, one can evaluate the degree of synergy of a specific model and easily understand the differences in the results. A strong synergy should be concluded if the observed response goes beyond the expected responses of all the four models.

### Prioritization of drug combinations

In addition to synergy, the sensitivity of a drug combination is equally important [8]. Prioritization of drug combinations based on the degree of synergy may only result in an excessive number of false positives, as a clinical endpoint for approving a drug combination is commonly its efficacy rather than the synergy. In cases where a strong synergy is obtained, the drug combination may not necessarily reach sufficient levels of therapeutic efficacy. To capture the sensitivity of a drug combination, we previously developed a combination sensitivity score (CSS) model that calculates the relative inhibition of a drug combination based on the area under the log10 scaled dose-response curves at the IC50 doses of the constituent drugs [8]. The CSS has the same unit as a drug response (*i.e*., % inhibition), which makes it directly comparable with the synergy scores defined in Equations (1–4). Therefore, we propose a scatter plot of CSS versus synergy score for a set of drug combinations. The proposed sensitivity-synergy (SS) plot significantly aids in the prioritization of drug combinations from a high-throughput experiment.

### Annotation of drug combinations

Drugs and cell lines in a drug screening dataset are annotated by querying related databases. PubChem CID, Isomeric SMILES, and standard InChIKey for drugs are extracted from PubChem [20]. Furthermore, we provide the molecular formula, clinical stage, disease indication, cross references to multiple major chemical compound databases (*i.e*., ChEMBL [21], ChEBI [22], DrugBank [23], BindingDB [24], PharmGKB [25], Guide to Pharmacology [26], Selleck, and ZINC [27]), and drug target profiles from MICHA [28]. Cell line annotation is obtained from Cellosaurus [29] (Version 37.0).

## Results

### Analysis of three-drug combinations

To test the algorithms designed for higher-order combinations, we utilize a recent anti-malaria study in which 16 triple artemisinin-based combination therapies have been tested against 15 parasite lines [30]. The study aimed to identify synergistic partner drugs that can be combined with artemisinin to overcome the emergence of malaria resistance. For each triple drug combination, a 10 × 10 × 12 multi-dimensional dose matrix was constructed, resulting in n = 1200 viability values. The datasets are obtained from the NCATS data portal at https://tripod.nih.gov/matrix-client/.

We analyze the triple drug combinations in terms of their sensitivity and synergy and visualize them in surface plots using dimension reduction techniques. **Figure 4** shows a synergistic triple combination, pyronaridine tetraphosphate-piperaquine-darunavir ethanolate (PYR-PQP-DRV) compared to an antagonistic combination, pyronaridine tetraphosphate-piperaquine-lopinavir (PYR-PQP-LPV). Despite similar sensitivities (the average % inhibition is 78.87 for PYR-PQP-DRV and 74.97 for PYR-PQP-LPV), their synergy scores differ drastically (the average ZIP score is 12.02 for PYR-PQP-DRV and −13.39 for PYR-PQP-LPV), suggesting more complex behaviors of the drug interactions as the number of drugs increases. Because of the lack of computational tools for analyzing higher-order drug combinations, the authors of the anti-malaria studies provided only the results of pairwise drug combinations, whereas the actual interactions of the three drugs were unexplored. Therefore, our method may provide novel insights that leverage higher-order drug combination design in a more tailored manner.

**Figure 4.**
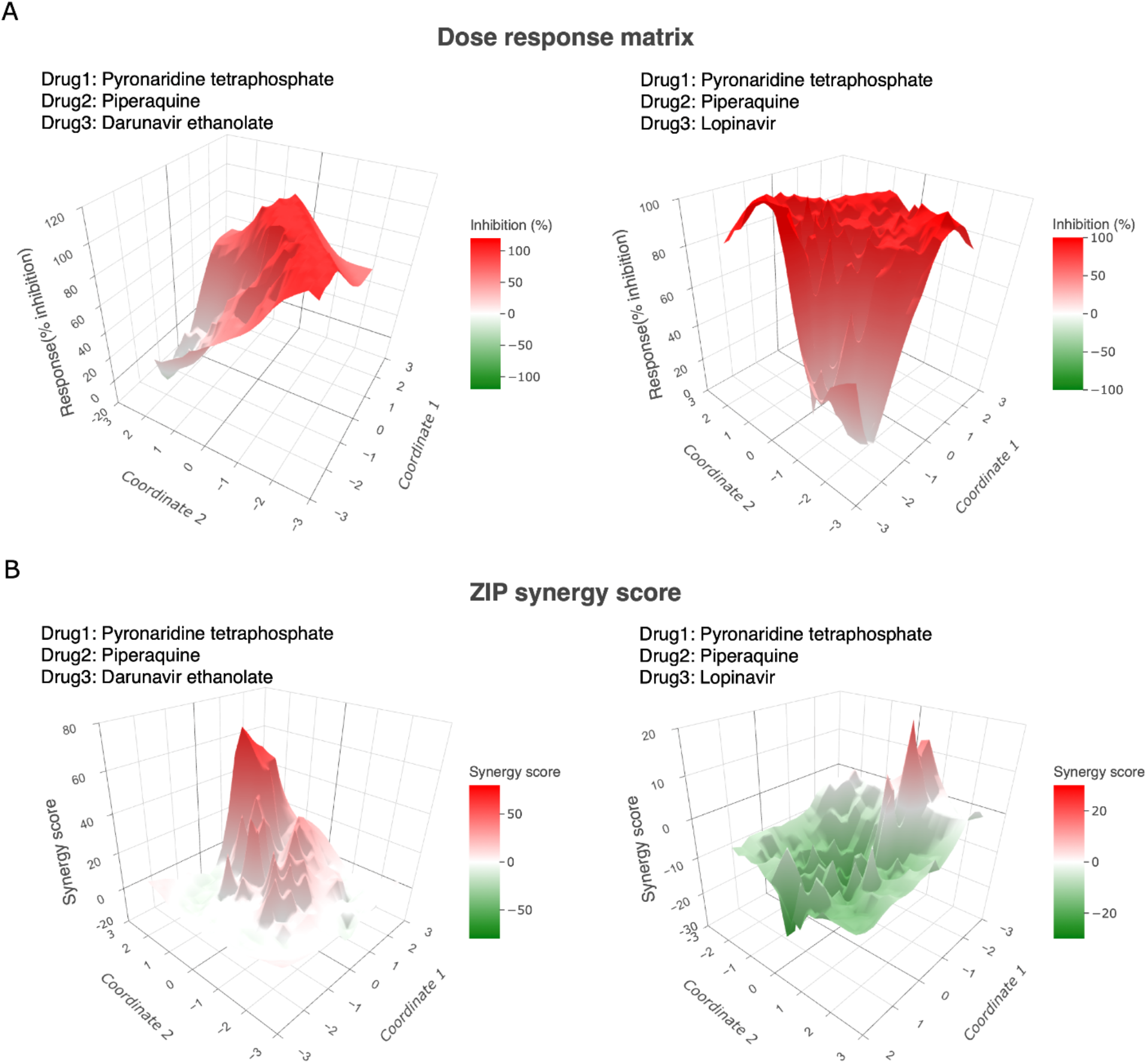
Example of surface plots for higher-order drug combinations. **A**. The dose-response map for a synergistic drug combination (pyronaridine tetraphosphate-piperaquine-darunavir ethanolate, left panel) compared to an antagonistic drug combination (pyronaridine tetraphosphate-piperaquine-lopinavir, right panel). **B**. The ZIP synergy score map for the same drug combinations. For surface plots of other synergy scores, including HSA, BLISS, and LOEWE, see Figure S2.

### Analysis of two-drug combinations with replicates

We obtain the ONEIL dataset from the DrugComb data portal [31]. The ONEIL data is a pan-cancer drug combination study in which 583 drug combinations were tested across 39 cell lines [32]. Four replicates were produced for each drug combination, making it a representative example dataset to analyze the significance of the synergy scores. In **Figure 5**, we show the selective drug combinations tested in the A2058 cell line (melanoma).

**Figure 5.**
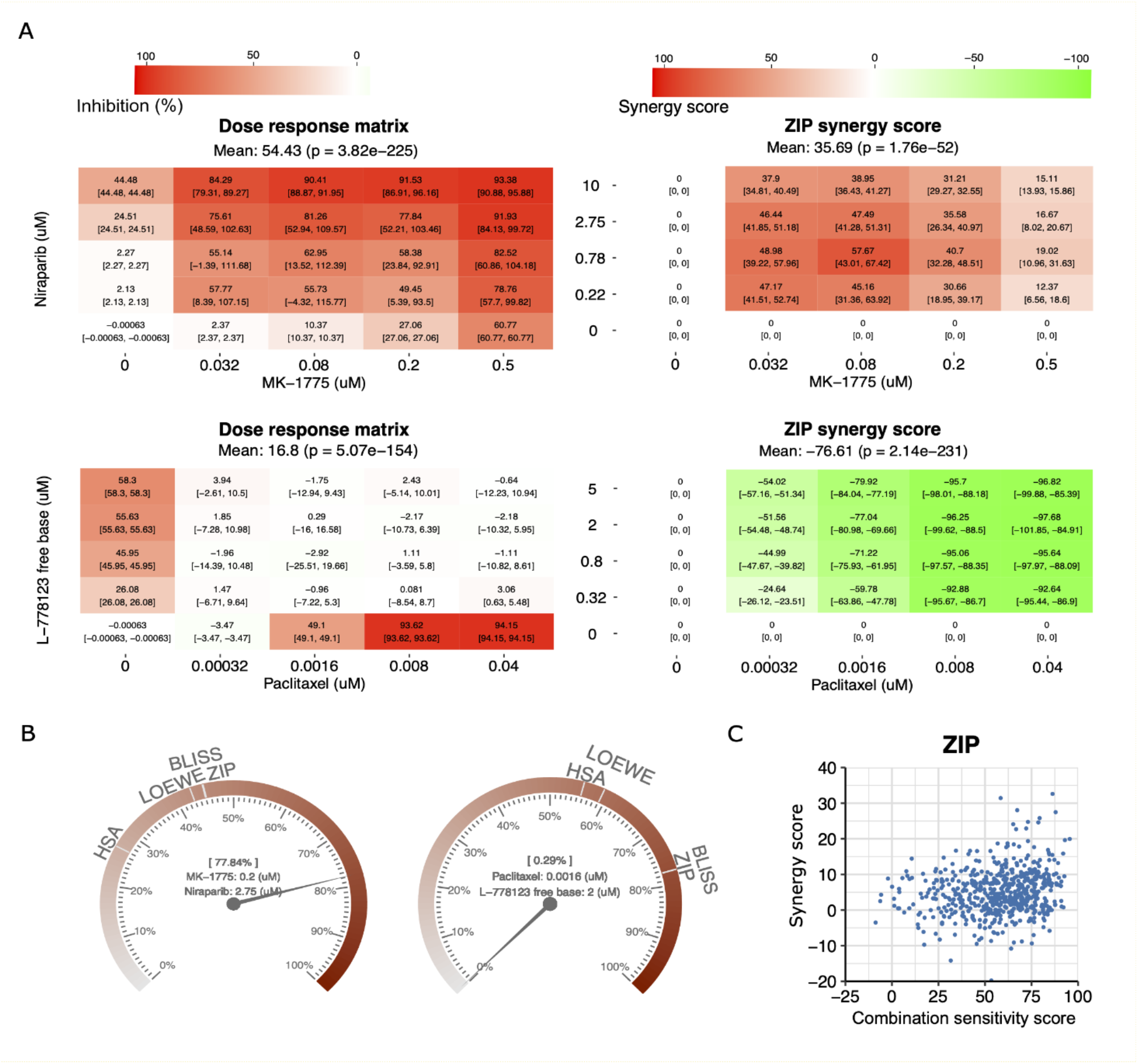
Example of visualizations for two-drug combinations with replicates. **A**. An example of the confidence interval plots of a synergistic drug pair (MK-1775 and niraparib; upper panel) and an antagonistic drug pair (paclitaxel and L-778123; lower panel). **B**. The synergy barometer for a given dose combination (left panel: MK-1775 at 0.2 μM in combination with niraparib at 2.75 μM; right panel: paclitaxel at 0.0016 μM in combination with L-778123 at 2 μM). **C**. The synergy-sensitivity plot for drug pairs tested in the A2058 cell line.

As shown in Figure 5A, the confidence intervals for the responses and synergy scores vary at different doses, suggesting that not all scores are equally significant. Furthermore, we observe a general trend that a higher response or synergy score tends to have a smaller confidence interval, consistent with the expectation of a typical drug screening experiment. To evaluate the synergy scores more systematically, we provide a synergy barometer for specific dose combinations. For example, for MK-1775 at 0.2 μM combined with niraparib at 2.75 μM, the drug combination response reaches 77.84% inhibition, as indicated by the pointer readout on the barometer (Figure 5B, left panel). Such a drug combination response is synergistic, as the expected responses of the HSA, LOEWE, BLISS, and ZIP models are much smaller, shown as the marks on the barometer. In contrast, paclitaxel at 0.0016 μM in combination with L-778123 at 2 μM shows a near-zero response (0.29%inhibition), which is much smaller than the expected response among all the synergy models (Figure 5B, right panel); therefore, it is considered strong antagonism. Using the synergy barometer, the actual drug combination response can be directly compared with the expectations of non-interaction among multiple synergy models.

Furthermore, we also show the CSS and ZIP synergy scores of all the drug combinations tested in the A2058 cell line in an SS plot (Figure 5C, Figure S3). CSS indicates the efficacy of a drug combination, whereas the synergy score indicates the degree of interaction. To prioritize potential drug combinations, it is necessary to identify those with higher CSS and higher synergy scores, that is, the top-right corner of the SS plot.

### Annotation of mechanisms of action of drug combinations

Recent advances in machine learning and artificial intelligence studies have shown great potential in predictive modeling of drug synergies [33–35]. In these studies, the chemical structure information regarding drugs and the molecular profiles of cells are considered predictive features. However, for most of the drug combination datasets, annotations of drug and cell features are not provided. To facilitate the interpretation of mechanisms of action of drug combinations, it would be beneficial to provide data integration tools for effective annotation and accurate matching to public databases such that more comprehensive features used for synergy predictions can be obtained. Efforts to annotate and harmonize existing drug screening data, such as DrugComb [31,36] and DrugCombDB [37], have significantly contributed to the development of data-driven pharmacological modeling [38–40]. For newly generated drug screening datasets, we develop the SynergyFinder Plus portal as a companion tool for retrieving publicly available information in a more automated fashion. With one click of the button, users can obtain cross-database identifiers of the drugs and cells, as well as additional information, such as the mechanism of action and disease indication (**Figure 6**).

**Figure 6.**
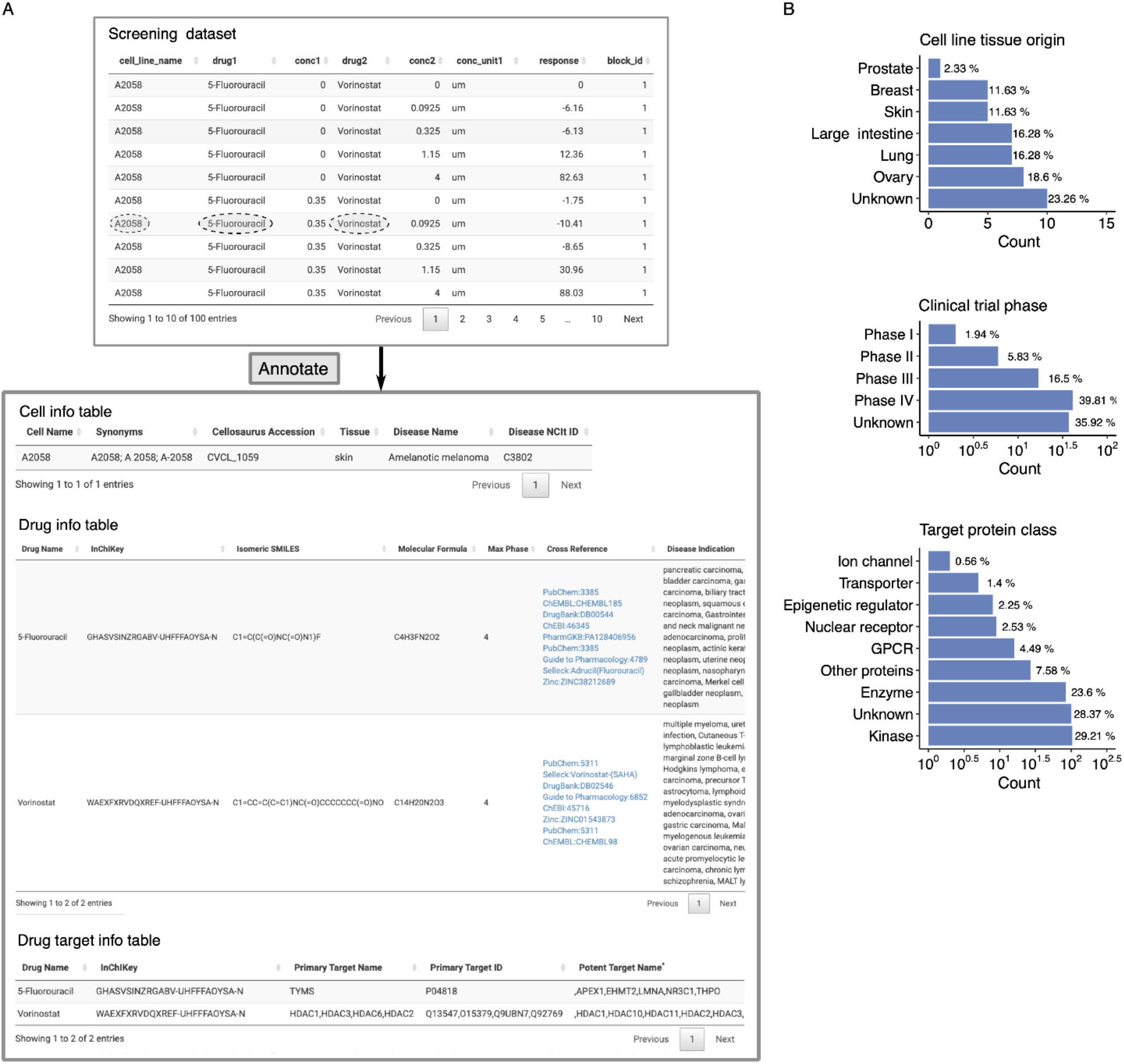
Example of mechanisms of action annotation for drug combination. **A**. An example of a drug screening annotation table. The tested drug combination, 5-fluorouracil and vorinostat, is further annotated with their chemical and target information. Cell line information for the A2058 is also provided, including synonyms, tissue, and disease types. **B**. Summary of drug clinical stage, drug target protein class, and cell line tissue origin for the ONEIL dataset.

We show how SynergyFinder Plus annotates a particular drug combination in the ONEIL [32] dataset (5-fluorouracil and vorinostat tested in the A2058 cell line, Figure 6A). After users upload the dataset, SynergyFinder Plus returns two annotation tables describing the cell and drug identities. For example, the cell annotation table includes synonyms, Cellosaurus accession, tissue origin, disease name, and disease NCI Thesaurus ID, whereas the drug annotation table includes InChIKey, isomeric SMILES, molecular formula, clinical trial phase, cross references, and disease indication. In addition, a third table listing primary and potent drug target information is provided. These annotations enable a more systematic exploration of drugs and cell lines tested in a high-throughput experiment (Figure 6B). With such a comprehensive collection of identifiers, we believe that the drug combination data can be better integrated with other related data sources for further downstream analysis.

## Discussion

Synergistic and effective drug combinations have long been studied to improve disease management. Testing potential drug combinations using clinically representative cell cultures has become an important tool for detecting signs of drug sensitivity and resistance; in some cases, the results have already provided informative decision-making resources to guide patient stratification in clinical trials [41]. To facilitate the prioritization of drug combinations, informatics tools that enable formal assessment of the degree of synergy as well as sensitivity are highly needed. We provide a major update to the commonly used SynergyFinder R package, which models drug interactions for any higher-order combinations [15]. Higher-order drug combinations have been recently explored in lung cancer [42], colorectal cancer [43], and ovarian cancer [44]. Therefore, it is expected that promising multi-drug combinations will enter formal clinical trials for multiple diseases [45]. Furthermore, we develope a statistical analysis of drug combinations, which is also generally applicable for higher-order combinations. Importantly, the mathematical models for higher-order combinations have been consistent with those for two-drug combinations; therefore, their scores are readily comparable across different orders. To visualize higher-order combinations, we develop a dimension reduction technique to display the synergy and the sensitivity landscape on a two-dimensional plane.

In addition to providing multiple synergy models, we address the problem of harmonization. With the increasing number of models that have been proposed to characterize drug synergy, there is an increasing need to help users understand the differences between these models such that a user may make an informed interpretation of the drug combination results. We propose the use of a synergy barometer on which all the synergy models can be systematically compared with the observed drug combination response. Despite only including the four major synergy models, HSA, BLISS, LOEWE, and ZIP, we propose the incorporation of other synergy models that can readily have their synergy scores put into the barometer for comparison with existing models. With the synergy barometer, we hope that the drug discovery community may derive a consensus on the best reporting guideline for drug combination data analysis, which has been long sought since the time of the Saariselkä agreement [46] and thereafter [47].

We also provide the formal statistical analysis for replicates of drug combinations. The confidence intervals and *P* values for all four synergy scores are comparable. Even for the cases in which no replicates have been performed, the significance of synergy over all the dose conditions can be provided, thus enabling a statistical evaluation of drug combinations in a primary screen that usually does not provide replicates due to the high number of combinations. Once a drug combination hit passes the initial screening based on the synergy barometer and the statistical significance, we recommend a confirmation screen that involves more doses and replicates to formally evaluate its synergy. If a user would like to study the mechanism of action of the drug combination, we also provide integrative tools to retrieve the chemical structure information and other functional annotations of the drugs, which will help in downstream analysis of the drug combinations. For higher-order drug combinations, the interpretation of drug synergies is a challenging problem. We hope that our tool provides the first evidence to pinpoint synergistic higher-order drug combinations.

Finally, we provide new features to harmonize the assessment of synergy and sensitivity. The synergy models that have been developed in the SynergyFinder R package have the same scale as the drug combination sensitivity method that was developed earlier, thus allowing a direct comparison in an SS plot. We recommend that users consider the SS plot when reporting their drug combination results, as sensitivity is the clinical endpoint for approving any drug or drug combinations. The drug combination sensitivity values also have the same scale as monotherapy drug responses in the unit of percentage inhibition; therefore, we can readily compare a drug combination with a single drug to evaluate their efficacy. With the harmonization of drug combination sensitivity and monotherapy sensitivity, we may provide an integrated data source for developing advanced machine learning approaches for predicting drug sensitivity, such as drug sensitivity datasets deposited in DrugComb [31]. To facilitate the FAIRification (findable, accessible, interoperable, and reusable characteristics) of drug combination analysis, we provide a web server at www.synergyfinderplus.org to allow users to upload the data and obtain the analysis results in the form of reports. In addition, we encourage users to deposit their experimental data in DrugComb at https://drugcomb.org/, currently one of the most comprehensive drug screening databases. With the algorithms and the source code made available through the SynergyFinder R package, we welcome the machine learning community to leverage the harmonized drug combination and monotherapy datasets for more robust and transferable predictions to further facilitate the drug combination discovery [34,48–50].

## Availability

The www.synergyfinderplus.org web server is located at the CSC-IT Center for Science in Finland. All the source code for implementing the data analysis and visualization is available as the SynergyFinder R package at https://bioconductor.org/packages/release/bioc/html/synergyfinder.html.

## Author CrediT statement

**Shuyu Zheng**: Conceptualization, Methodology, Software, Writing – Original draft preparation **Wenyu Wang**: Visualization, Writing – Original draft preparation **Jehad Aldahdooh**: Software **Alina Malyutina**: Methodology **Tolou Shadbahr**: Data Curation **Ziaurrehman Tanoli**: Resources **Alberto Pessia**: Methodology, Validation, Writing – Review & Editing **Jing Tang**: Supervision, Conceptualization, Methodology, Software, Writing – Original draft preparation

## Acknowledgments

This work is supported by the European Research Council (ERC) starting grant DrugComb (Informatics approaches for the rational selection of personalized cancer drug combinations) [No. 716063], European Commission H2020 EOSC-life (Providing an open collaborative space for digital biology in Europe [No. 824087], and the Academy of Finland grant [No. 317680]. The Sigrid Jusélius Foundation grant. SZ and WW hold a salaried position funded by the University of Helsinki through the Doctoral Program of Biomedicine (DPBM). WW also receives a personal grant from K. Albin Johanssons stiftelse. AM holds a salaried position funded by the University of Helsinki through the Doctoral Program of Integrative Life Science (ILS), and receives a personal grant from the Cancer Foundation Finland.

## Conflict of interests

None declared.

## Supplementary material

**Figure S1.**
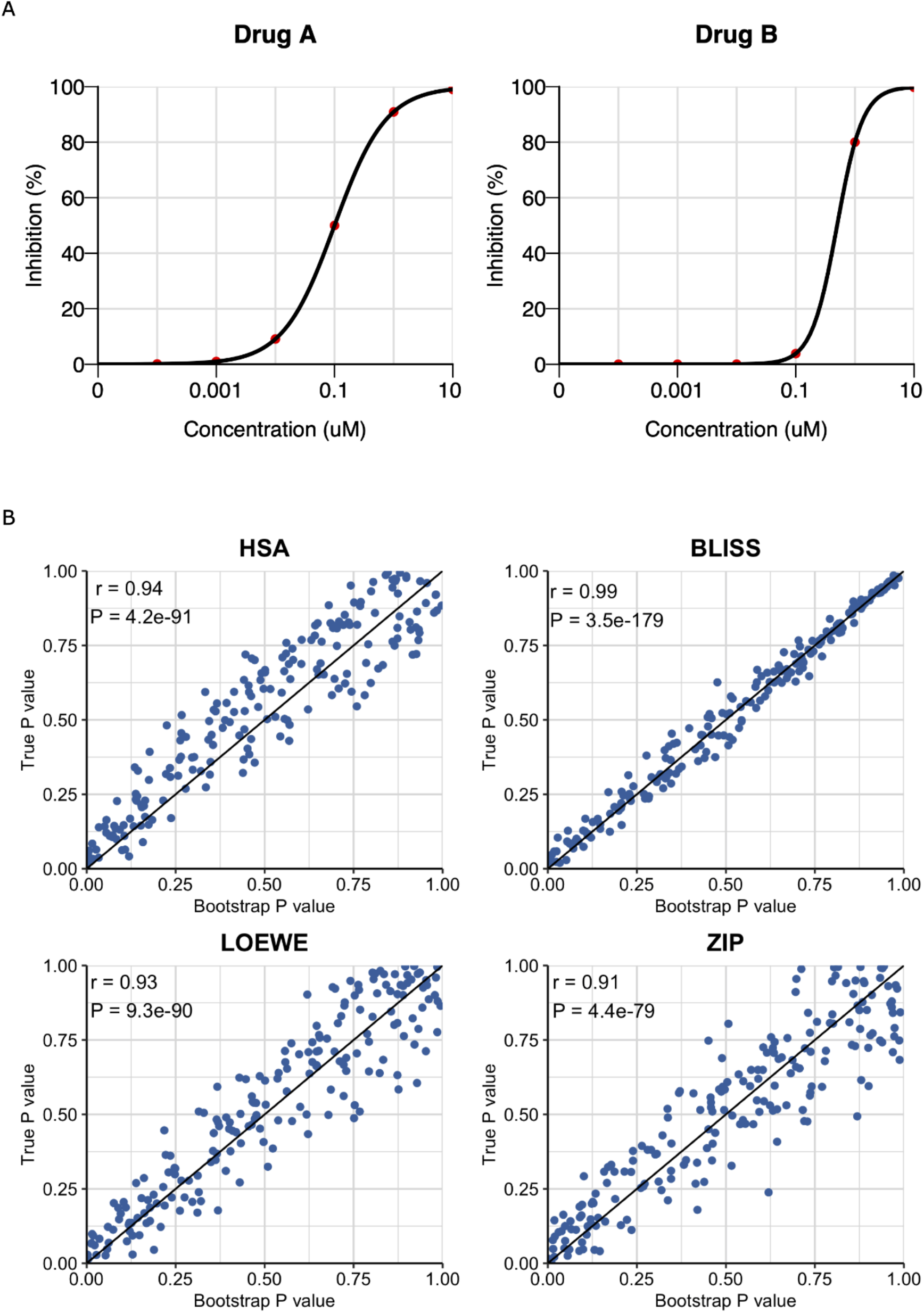
Accuracy of *P* values for synergy scores by bootstrapping. **A.** The expected monotherapy responses are defined by four-parameter log-logistic functions and the expected combinational responses are defined by the non-interaction models. Based on the expected values, the distribution of the test statistic under null hypothesis can be determined by simulations. **B**. The scatter plots for comparing the true *P* values and bootstrapping *P* values for 200 samples.

**Figure S2.**
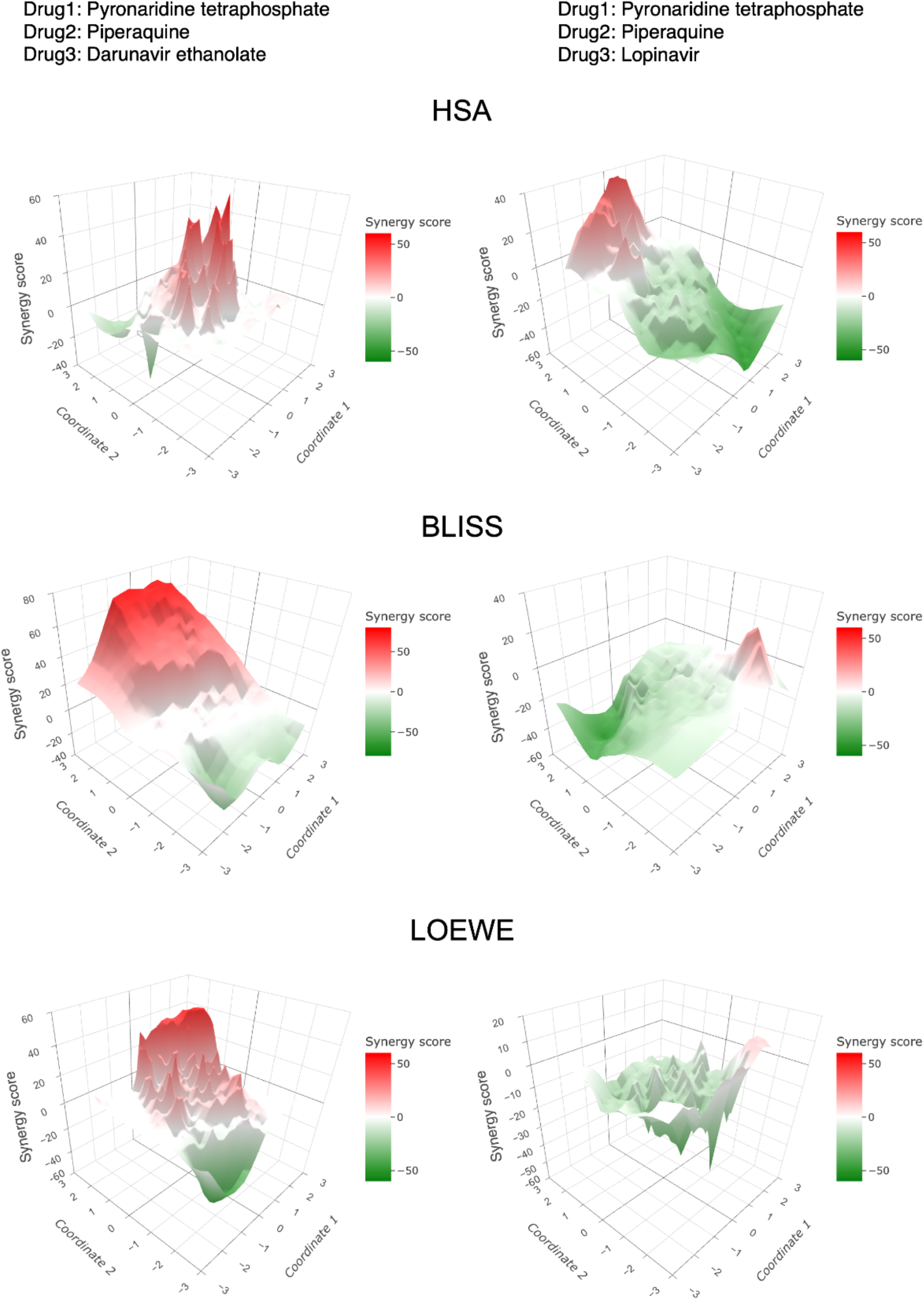
Two-dimensional surface plots of synergy scores. **A.** HSA, **B.** BLISS, and **C.** LOEWE for a synergistic three-drug combination (pyronaridine tetraphosphate-piperaquine-darunavir ethanolate, left panel) compared to an antagonistic drug combination (pyronaridine tetraphosphate-piperaquine-lopinavir, right panel).

**Figure S3.**
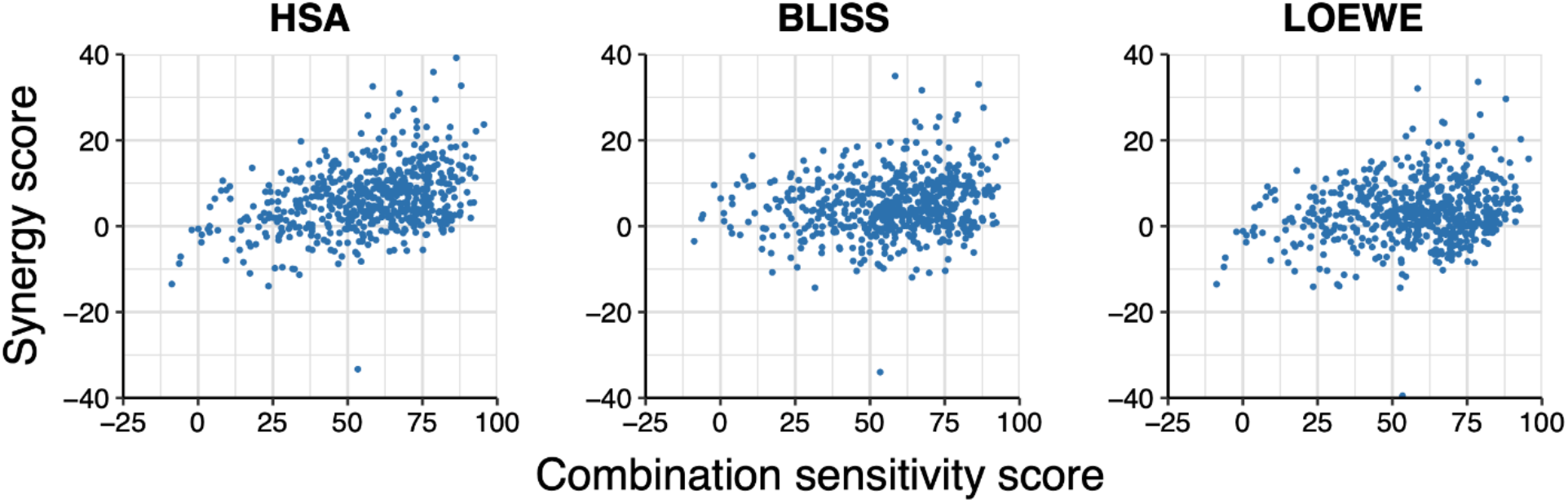
Synergy-sensitivity plots for HSA, BLISS, and LOEWE scores in relation to the combination sensitivity score for the drug pairs tested in the A2058 cell line.

**Table S1.**
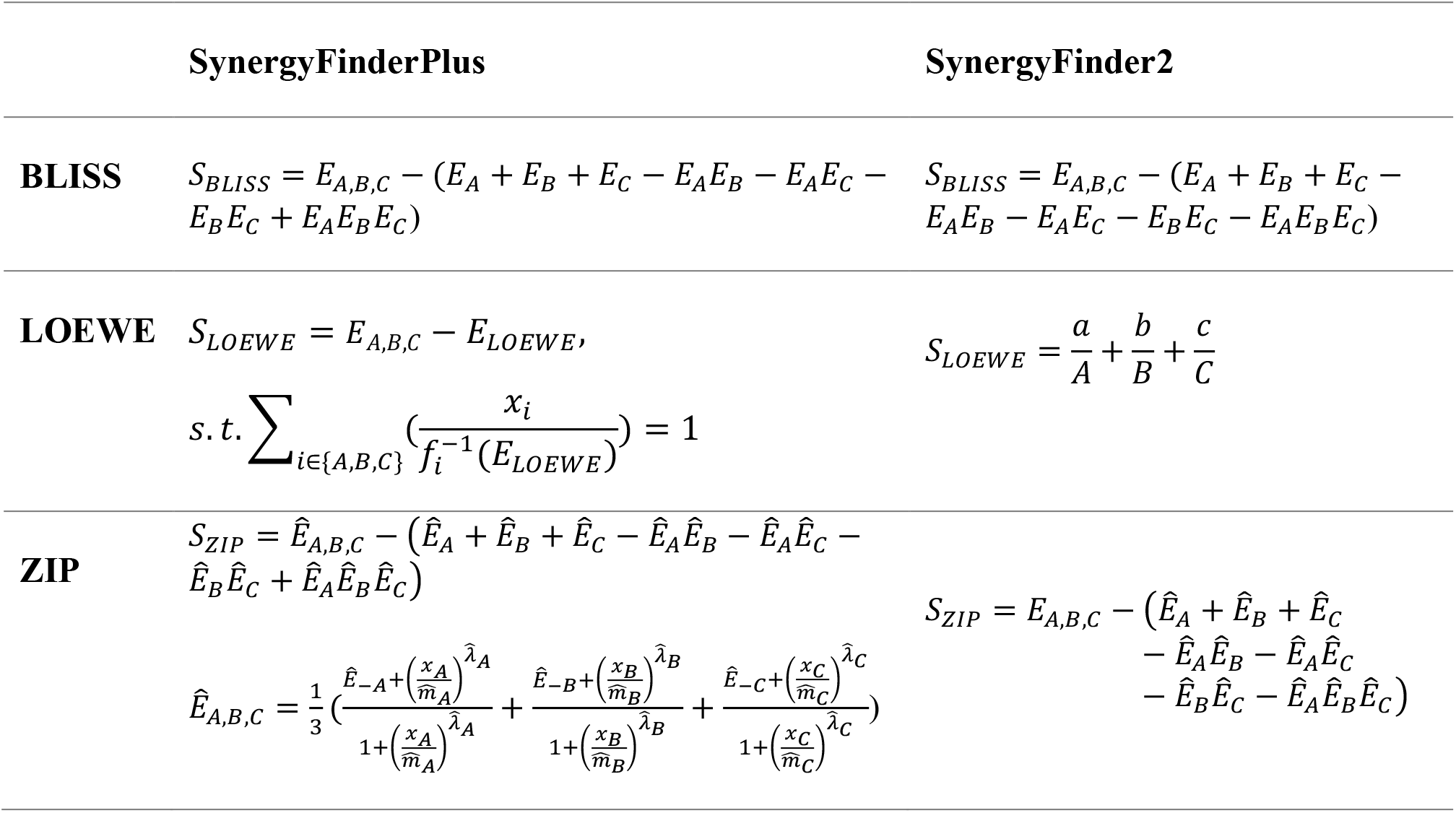
The key differences in the mathematical modeling between SynergyFinderPlus and SynergyFinder2 for three-drug combinations.

|  | SynergyFinderPlus | SynergyFinder2 |
| --- | --- | --- |
| **BLISS** | $S_{BLISS} = E_{A,B,C} - (E_A + E_B + E_C - E_A E_B - E_A E_C - E_B E_C + E_A E_B E_C)$ | $S_{BLISS} = E_{A,B,C} - (E_A + E_B + E_C - E_A E_B - E_A E_C - E_B E_C - E_A E_B E_C)$ |
| **LOEWE** | $S_{LOEWE} = E_{A,B,C} - E_{LOEWE},$<br>$s.t. \sum_{i \in \{A,B,C\}} (\frac{x_i}{f_i^{-1}(E_{LOEWE})}) = 1$ | $S_{LOEWE} = \frac{a}{A} + \frac{b}{B} + \frac{c}{C}$ |
| **ZIP** | $S_{ZIP} = \hat{E}_{A,B,C} - (\hat{E}_A + \hat{E}_B + \hat{E}_C - \hat{E}_A \hat{E}_B - \hat{E}_A \hat{E}_C - \hat{E}_B \hat{E}_C + \hat{E}_A \hat{E}_B \hat{E}_C)$<br>$\hat{E}_{A,B,C} = \frac{1}{3} (\frac{\hat{E}_{-A} + (\frac{x_A}{\hat{m}_A})^{\hat{\lambda}_A}}{1 + (\frac{x_A}{\hat{m}_A})^{\hat{\lambda}_A}} + \frac{\hat{E}_{-B} + (\frac{x_B}{\hat{m}_B})^{\hat{\lambda}_B}}{1 + (\frac{x_B}{\hat{m}_B})^{\hat{\lambda}_B}} + \frac{\hat{E}_{-C} + (\frac{x_C}{\hat{m}_C})^{\hat{\lambda}_C}}{1 + (\frac{x_C}{\hat{m}_C})^{\hat{\lambda}_C}})$ | $S_{ZIP} = E_{A,B,C} - (\hat{E}_A + \hat{E}_B + \hat{E}_C - \hat{E}_A \hat{E}_B - \hat{E}_A \hat{E}_C - \hat{E}_B \hat{E}_C - \hat{E}_A \hat{E}_B \hat{E}_C)$ |

**File S1 The simulation code for testing accuracy of *P* values for synergy scores by bootstrapping**

## Notes

http://www.synergyfinderplus.org

### Competing Interest Statement

The authors have declared no competing interest.

### Summary of Updates

1. Title is changed a little bit to fit the format required by the journal. 2. Ziaurrehman Tanoli is added to the author list. 3. Section "Visualization of higher-order synergy and sensitivity scores" in "Methods" is updated to clarify the dimension reduction method. 4. Figure 3 and Section "Statistical analysis" in the "Methods" is updated to clarify the bootstrap methods we are using for statistical analysis. Figure S1 is added to clarify the accuracy of the P values from bootstrap. 5. File S1 is added. It is the R code for testing the bootstrapping P values. 6. Figure 5, Figure 6, and Figure S3 (Figure S2 in the previous submission) are updated to show the updated results. 7. Figure 1, Figure 2, Figure 4, and Figure S2 (Figure S1 in the previous submission) are updated to fit the format required by the journal. 8. Table S1 is added to clarify the differences between SynergyFinder Plus and SynergyFinder2. 9. Seven references (21 - 27) are added to clarify the resources of drug annotation data.

