## Supplementary material for "SynergyFinder Plus: Toward Better Interpretation and Annotation of Drug Combination Screening Datasets": File S1

R scrip for testing accuracy of *P* values for synergy scores by bootstrapping

\# Install SnergyFinder R package ------------------------------------------

\# SynergyFinder is published on Bioconductor

\# https://www.bioconductor.org/packages/release/bioc/html/synergyfinder.html

\#

\# if (!requireNamespace("BiocManager", quietly = TRUE))

\#	install.packages("BiocManager")

\#

\# BiocManager::install("synergyfinder")

\# Load packages -----------------------------------------------------------

library(synergyfinder)

library(ggplot2)

\# Define functions --------------------------------------------------------

\# 4-parameter log-logistic function

f_ll <- function(x, p) {

p[1] + (p[2] - p[1]) * x^p[3] / (x^p[3] + p[4]^p[3])

}

\# Transform the P value format in the bootstrap result

translate_p_value <- function(s){

if (grepl(s, "e", fixed = T)){

s <- gsub("<", "", s)

s <- unlist(strsplit(s, "e"))

}

n <- as.numeric(s[1]) * 10 ^ as.numeric(s[2])

return(n)

}

\# Define variables --------------------------------------------------------

\# Parameters for mono-therapy dose-response curve

p_a <- c("lower" = 0, "upper" = 1, "hill" = 1, "ic50" = 0.1)

p_b <- c("lower" = 0, "upper" = 1, "hill" = 2, "ic50" = 0.5)

\# Number of replications per dose

n <- 3

\# Vector of tested doses

dose <- c(0.0001, 0.001, 0.01, 0.1, 1, 10)

k <- length(dose)

\# Combination doses

dose_comb <- as.matrix(expand.grid(dose, dose))

v <- nrow(dose_comb)

\# Constant standard deviation for each dose combination

s <- 0.05

\# Number of bootstraps

n_bootstrap <- 10000

\# Number of simulations

N <- n_bootstrap

\# Bliss score -------------------------------------------------------------

\# Output directory

output_dir <- paste0("../data/Bliss_synergyfinder_", n_bootstrap, "/")

if (!dir.exists(output_dir)){

dir.create(output_dir, recursive = TRUE)

}

\## Expected response values -----------------------------------------------

\# The mono-therapy expected values (from log-logistic dose-response curve)

m_a <- rep(f_ll(dose, p_a), each = n)

m_b <- rep(f_ll(dose, p_b), each = n)

\# Bliss model for inhibition (hill coefficient positive)

f_bliss <- function(x, p_a, p_b) {

f_ll(x[1], p_a) + f_ll(x[2], p_b) - f_ll(x[1], p_a) * f_ll(x[2], p_b)

}

\# The combo-therapy expected values (H0: bliss model)

m_ab <- rep(apply(dose_comb, 1, f_bliss, p_a, p_b), each = n)

\## Distribution under null hypothesis -------------------------------------

\# Simulating the distribution of the test statistic under the null hypothesis

t_sample <- rep(0, N)

for (i in seq_len(N)) {

\# simulate mono-therapy data

y_a <- rnorm(n = length(m_a), mean = m_a, sd = s)

y_b <- rnorm(n = length(m_b), mean = m_b, sd = s)

\# simulate combo-therapy data

y_ab <- rnorm(n = length(m_ab), mean = m_ab, sd = s)

\# compute the sample means for each dose/combination

obs_m_a <- c(by(y_a, rep(seq_len(k), each = n), mean))

obs_m_b <- c(by(y_b, rep(seq_len(k), each = n), mean))

obs_m_ab <- c(by(y_ab, rep(seq_len(v), each = n), mean))

\# cross the previous sample means to compute the bliss value

tmp <- as.matrix(expand.grid(obs_m_a, obs_m_b))

obs_bliss <- tmp[, 1] + tmp[, 2] - tmp[, 1] * tmp[, 2]

\# synergy score

synsco <- obs_m_ab - obs_bliss

\# average of the synergy score over the whole matrix

t_sample[i] <- mean(synsco)

}

\## Bootstrap -------------------------------------------------------------

\# Bootstrap function for Bliss model

bootstrap <- function(i, m_a, m_b, m_ab, s, N, dose_comb, k, t_sample){

set.seed(i)

\# generate a new sample for the bootstrap study

y_a <- rnorm(n = length(m_a), mean = m_a, sd = s)

y_b <- rnorm(n = length(m_b), mean = m_b, sd = s)

y_ab <- rnorm(n = length(m_ab), mean = m_ab, sd = s)

\# compute the observed statistics

obs_m_a <- c(by(y_a, rep(seq_len(k), each = n), mean))

obs_m_b <- c(by(y_b, rep(seq_len(k), each = n), mean))

obs_m_ab <- c(by(y_ab, rep(seq_len(v), each = n), mean))

response <- data.frame(

conc1 = c(0, dose, rep(0, k), dose_comb[, "Var1"]),

conc2 = c(0, rep(0, k), dose, dose_comb[, "Var2"]),

response = c(0, obs_m_a, obs_m_b, obs_m_ab) * 100,

stringsAsFactors = FALSE

)

obs_Bliss <- synergyfinder::Bliss(response)

t_obs <- mean(obs_Bliss$Bliss_synergy[(2*k+2):nrow(obs_Bliss)])/100

\# true p-value

true_pv <- mean((t_sample <= -abs(t_obs)) | (t_sample > abs(t_obs)))

\# do bootstrap...

n_well <- (1+2*k+nrow(dose_comb)) * n

data_table <- data.frame(

block_id = rep(1, times = n_well),

drug1 = rep("drug_a", times = n_well),

drug2 = rep("drug_b", times = n_well),

conc1 = rep(c(0, dose, rep(0, k), dose_comb[, "Var1"]), each = n),

conc2 = rep(c(0, rep(0, k), dose, dose_comb[, "Var2"]), each = n),

ConcUnit = rep("uM", times = n_well),

response = c(rep(0, times = n), y_a, y_b, y_ab) * 100,

stringsAsFactors = FALSE

)

data <- synergyfinder::ReshapeData(data_table, data_type = "inhibition")

res <- synergyfinder::CalculateSynergy(

data,

iteration = N,

seed = i,

method = "Bliss"

)

boot_pv <- res$drug_pairs

boot_pv$iteration = i

boot_pv$true_pv <- true_pv

\# plot(density(t_sample), xlim = c(-0.05, 0.05))

\# plot(density(t_boot), xlim = c(-0.05, 0.05))

return(

p_values = boot_pv

)

}

\# Run bootstrap

result <- NULL

for (i in 1:n_bootstrap) {

tmp <- bootstrap(

job_id, m_a, m_b, m_ab, s,

N = n_bootstrap, dose_comb, k, t_sample

)

result <- rbind.data.frame(result, tmp)

}

if (!is.numeric(result$Bliss_synergy_p_value)){

result$Bliss_synergy_p_value <- sapply (

result$Bliss_synergy_p_value,

translate_p_value

)

}

write.csv(result, paste0(output_dir, "P_value.csv"), row.names = FALSE)

\## Visualization ----------------------------------------------------------

c <- cor.test(result$Bliss_synergy_p_value, result$true_pv)

p <- ggplot(result, aes(x = Bliss_synergy_p_value, y = true_pv)) +

geom_point(color = "#2D72AD") +

labs(

x = "Bootstrap P value",

y = "True P value",

title = "Bliss"

) +

geom_abline(intercept = 0, slope = 1) +

theme_classic()+

annotate(

geom = "text",

x = min(result$Bliss_synergy_p_value),

y = max(result$true_pv) - 0.1,

label = paste0(

"r = ", signif(c$estimate, 2),

"\nP = ", signif(c$p.value, 2)

),

hjust = 0

) +

theme(

panel.background = ggplot2::element_rect(

fill = "white",

colour = "white",

size = 2,

linetype = "solid"

),

panel.grid.major = ggplot2::element_line(

size = 0.5,

linetype = 'solid',

colour = "#DFDFDF"

),

panel.grid.minor = ggplot2::element_line(

size = 0.25,

linetype = 'solid',

colour = "#DFDFDF"

),

plot.title = ggplot2::element_text(

size = 13.5,

face = "bold",

hjust = 0.5

),

axis.text = ggplot2::element_text(

size = 10,

color = "black"

),

axis.title = ggplot2::element_text(

size = 10,

face = "italic"

),

\# axis.ticks = element_blank(),

axis.line.y.left = ggplot2::element_line(color = "black"),

axis.line.x.bottom = ggplot2::element_line(color = "black")

)

ggsave(

paste0(output_dir, "Scatter_plot_for_p_values.png"),

p, width = 3.5, height = 3

)

\# HSA score ---------------------------------------------------------------

\# Clean the environment

common_variables <- c(

"p_a", "p_b", "n", "dose", "k", "dose_comb",

"v", "s", "n_bootstrap", "N", "f_ll", "translate_p_value"

)

rm(list = setdiff(ls(), common_variables))

\# Output directory

output_dir <- paste0("../data/HSA_synergyfinder_", n_bootstrap, "/")

if (!dir.exists(output_dir)){

dir.create(output_dir, recursive = TRUE)

}

\## Expected response values -----------------------------------------------

\# The mono-therapy expected values (log-logistic dose-response curve)

m_a <- rep(f_ll(dose, p_a), each = n)

m_b <- rep(f_ll(dose, p_b), each = n)

\# HSA model for inhibition (hill coefficient positive)

f_hsa <- function(x, p_a, p_b){

max(f_ll(x[1], p_a), f_ll(x[2], p_b))

}

\# The combo-therapy expected values (H0: HSA model)

m_ab <- rep(apply(dose_comb, 1, f_hsa, p_a, p_b), each = n)

\## Distribution under null hypothesis -------------------------------------

\# Simulating the distribution of the test statistic under the null hypothesis

t_sample <- rep(0, N)

for (i in seq_len(N)) {

\# simulate mono-therapy data

y_a <- rnorm(n = length(m_a), mean = m_a, sd = s)

y_b <- rnorm(n = length(m_b), mean = m_b, sd = s)

\# simulate combo-therapy data

y_ab <- rnorm(n = length(m_ab), mean = m_ab, sd = s)

\# compute the sample means for each dose/combination

obs_m_a <- c(by(y_a, rep(seq_len(k), each = n), mean))

obs_m_b <- c(by(y_b, rep(seq_len(k), each = n), mean))

obs_m_ab <- c(by(y_ab, rep(seq_len(v), each = n), mean))

\# cross the previous sample means to compute the hsa value

tmp <- as.matrix(expand.grid(obs_m_a, obs_m_b))

obs_hsa <- pmax(tmp[, 1], tmp[, 2])

\# synergy score

synsco <- obs_m_ab - obs_hsa

\# average of the synergy score over the whole matrix

t_sample[i] <- mean(synsco)

}

\## Bootstrap --------------------------------------------------------------

\# Bootstrap function for HSA model

bootstrap <- function(i, m_a, m_b, m_ab, s, N, dose_comb, k, t_sample){

set.seed(i)

\# generate a new sample for the bootstrap study

y_a <- rnorm(n = length(m_a), mean = m_a, sd = s)

y_b <- rnorm(n = length(m_b), mean = m_b, sd = s)

y_ab <- rnorm(n = length(m_ab), mean = m_ab, sd = s)

\# compute the observed statistics

obs_m_a <- c(by(y_a, rep(seq_len(k), each = n), mean))

obs_m_b <- c(by(y_b, rep(seq_len(k), each = n), mean))

obs_m_ab <- c(by(y_ab, rep(seq_len(v), each = n), mean))

response <- data.frame(

conc1 = c(0, dose, rep(0, k), dose_comb[, "Var1"]),

conc2 = c(0, rep(0, k), dose, dose_comb[, "Var2"]),

response = c(0, obs_m_a, obs_m_b, obs_m_ab) * 100,

stringsAsFactors = FALSE

)

obs_HSA <- synergyfinder::HSA(response)

t_obs <- mean(obs_HSA$HSA_synergy[(2*k+2):nrow(obs_HSA)])/100

\# true p-value

true_pv <- mean((t_sample <= -abs(t_obs)) | (t_sample > abs(t_obs)))

\# do bootstrap...

n_well <- (1+2*k+nrow(dose_comb)) * n

data_table <- data.frame(

block_id = rep(1, times = n_well),

drug1 = rep("drug_a", times = n_well),

drug2 = rep("drug_b", times = n_well),

conc1 = rep(c(0, dose, rep(0, k), dose_comb[, "Var1"]), each = n),

conc2 = rep(c(0, rep(0, k), dose, dose_comb[, "Var2"]), each = n),

ConcUnit = rep("uM", times = n_well),

response = c(rep(0, times = n), y_a, y_b, y_ab) * 100,

stringsAsFactors = FALSE

)

data <- synergyfinder::ReshapeData(data_table, data_type = "inhibition")

res <- synergyfinder::CalculateSynergy(

data,

iteration = N,

seed = i,

method = "HSA"

)

boot_pv <- res$drug_pairs

boot_pv$iteration = i

boot_pv$true_pv <- true_pv

\# plot(density(t_sample), xlim = c(-0.05, 0.05))

\# plot(density(t_boot), xlim = c(-0.05, 0.05))

return(

p_values = boot_pv

)

}

\# Run bootstrap

result <- NULL

for (i in 1:n_bootstrap) {

tmp <- bootstrap(

job_id, m_a, m_b, m_ab, s,

N = n_bootstrap, dose_comb, k, t_sample

)

result <- rbind.data.frame(result, tmp)

}

if (!is.numeric(result$HSA_synergy_p_value)){

result$HSA_synergy_p_value <- sapply (

result$HSA_synergy_p_value,

translate_p_value

)

}

write.csv(result, paste0(output_dir, "P_value.csv"), row.names = FALSE)

\## Visualization ----------------------------------------------------------

c <- cor.test(result$HSA_synergy_p_value, result$true_pv)

p <- ggplot(result, aes(x = HSA_synergy_p_value, y = true_pv)) +

geom_point(color = "#2D72AD") +

labs(

x = "Bootstrap P value",

y = "True P value",

title = "HSA"

) +

geom_abline(intercept = 0, slope = 1) +

theme_classic()+

annotate(

geom = "text",

x = min(result$HSA_synergy_p_value),

y = max(result$true_pv) - 0.1,

label = paste0("r = ", signif(c$estimate, 2), "\nP = ", signif(c$p.value, 2)),

hjust = 0

) +

theme(

panel.background = ggplot2::element_rect(

fill = "white",

colour = "white",

size = 2,

linetype = "solid"

),

panel.grid.major = ggplot2::element_line(

size = 0.5,

linetype = 'solid',

colour = "#DFDFDF"

),

panel.grid.minor = ggplot2::element_line(

size = 0.25,

linetype = 'solid',

colour = "#DFDFDF"

),

plot.title = ggplot2::element_text(

size = 13.5,

face = "bold",

hjust = 0.5

),

axis.text = ggplot2::element_text(

size = 10,

color = "black"

),

axis.title = ggplot2::element_text(

size = 10,

face = "italic"

),

\# axis.ticks = element_blank(),

axis.line.y.left = ggplot2::element_line(color = "black"),

axis.line.x.bottom = ggplot2::element_line(color = "black")

)

ggsave(

paste0(output_dir, "Scatter_plot_for_p_values.png"),

p, width = 3.5, height = 3

)

\# ZIP score ---------------------------------------------------------------

\# Clean the environment

common_variables <- c(

"p_a", "p_b", "n", "dose", "k", "dose_comb",

"v", "s", "n_bootstrap", "N", "f_ll", "translate_p_value"

)

rm(list = setdiff(ls(), common_variables))

\# Output directory

output_dir <- paste0("../data/ZIP_synergyfinder_", n_bootstrap, "/")

if (!dir.exists(output_dir)){

dir.create(output_dir, recursive = TRUE)

}

\## Expected response values -----------------------------------------------

\# The mono-therapy expected values (log-logistic dose-response curve)

m_a <- rep(f_ll(dose, p_a), each = n)

m_b <- rep(f_ll(dose, p_b), each = n)

\# ZIP model for inhibition (hill coefficient positive)

f_zip <- function(x, p_a, p_b) {

f_ll(x[1], p_a) + f_ll(x[2], p_b) - f_ll(x[1], p_a) * f_ll(x[2], p_b)

}

\# The combo-therapy expected values (H0: ZIP model)

m_ab <- rep(apply(dose_comb, 1, f_zip, p_a, p_b), each = n)

\## Distribution under null hypothesis -------------------------------------

\# Simulating the distribution of the test statistic under the null hypothesis

t_sample <- rep(0, N)

for (i in seq_len(N)) {

\# simulate mono-therapy data

y_a <- rnorm(n = length(m_a), mean = m_a, sd = s)

y_b <- rnorm(n = length(m_b), mean = m_b, sd = s)

\# simulate combo-therapy data

y_ab <- rnorm(n = length(m_ab), mean = m_ab, sd = s)

\# compute the sample means for each dose/combination

obs_m_a <- c(by(y_a, rep(seq_len(k), each = n), mean))

obs_m_b <- c(by(y_b, rep(seq_len(k), each = n), mean))

obs_m_ab <- c(by(y_ab, rep(seq_len(v), each = n), mean))

\# cross the previous sample means to compute the zip value

tmp <- as.matrix(expand.grid(obs_m_a, obs_m_b))

concs <- as.matrix(expand.grid(obs_m_a, obs_m_b))

response <- data.frame(

conc1 = c(0, dose, rep(0, k), dose_comb[, "Var1"]),

conc2 = c(0, rep(0, k), dose, dose_comb[, "Var2"]),

response = c(0, obs_m_a, obs_m_b, obs_m_ab) * 100,

stringsAsFactors = FALSE

)

synsco <- synergyfinder::ZIP(response)

synsco <- mean(synsco$ZIP_synergy[(2*k+2):nrow(synsco)])/100

\# average of the synergy score over the whole matrix

t_sample[i] <- mean(synsco)

}

\## Bootstrap --------------------------------------------------------------

\# Bootstrap function for ZIP model

bootstrap <- function(i, m_a, m_b, m_ab, s, N, dose_comb, k, t_sample){

set.seed(i)

\# generate a new sample for the bootstrap study

y_a <- rnorm(n = length(m_a), mean = m_a, sd = s)

y_b <- rnorm(n = length(m_b), mean = m_b, sd = s)

y_ab <- rnorm(n = length(m_ab), mean = m_ab, sd = s)

\# compute the observed statistics

obs_m_a <- c(by(y_a, rep(seq_len(k), each = n), mean))

obs_m_b <- c(by(y_b, rep(seq_len(k), each = n), mean))

obs_m_ab <- c(by(y_ab, rep(seq_len(v), each = n), mean))

response <- data.frame(

conc1 = c(0, dose, rep(0, k), dose_comb[, "Var1"]),

conc2 = c(0, rep(0, k), dose, dose_comb[, "Var2"]),

response = c(0, obs_m_a, obs_m_b, obs_m_ab) * 100,

stringsAsFactors = FALSE

)

obs_zip <- synergyfinder::ZIP(response)

t_obs <- mean(obs_zip$ZIP_synergy[(2*k+2):nrow(obs_zip)])/100

\# true p-value

true_pv <- mean((t_sample <= -abs(t_obs)) | (t_sample > abs(t_obs)))

\# do bootstrap...

n_well <- (1+2*k+nrow(dose_comb)) * n

data_table <- data.frame(

block_id = rep(1, times = n_well),

drug1 = rep("drug_a", times = n_well),

drug2 = rep("drug_b", times = n_well),

conc1 = rep(c(0, dose, rep(0, k), dose_comb[, "Var1"]), each = n),

conc2 = rep(c(0, rep(0, k), dose, dose_comb[, "Var2"]), each = n),

ConcUnit = rep("uM", times = n_well),

response = c(rep(0, times = n), y_a, y_b, y_ab) * 100,

stringsAsFactors = FALSE

)

data <- synergyfinder::ReshapeData(data_table, data_type = "inhibition")

res <- synergyfinder::CalculateSynergy(

data,

iteration = N,

seed = i,

method = "ZIP"

)

boot_pv <- res$drug_pairs

boot_pv$iteration = i

boot_pv$true_pv <- true_pv

\# plot(density(t_sample), xlim = c(-0.05, 0.05))

\# plot(density(t_boot), xlim = c(-0.05, 0.05))

return(

p_values = boot_pv

)

}

\# Run bootstrap

result <- NULL

for (i in 1:n_bootstrap) {

tmp <- bootstrap(

job_id, m_a, m_b, m_ab, s,

N = n_bootstrap, dose_comb, k, t_sample

)

result <- rbind.data.frame(result, tmp)

}

if (!is.numeric(result$ZIP_synergy_p_value)){

result$ZIP_synergy_p_value <- sapply (

result$ZIP_synergy_p_value,

translate_p_value

)

}

write.csv(result, paste0(output_dir, "P_value.csv"), row.names = FALSE)

\## Visualization ----------------------------------------------------------

c <- cor.test(result$ZIP_synergy_p_value, result$true_pv)

p <- ggplot(result, aes(x = ZIP_synergy_p_value, y = true_pv)) +

geom_point(color = "#2D72AD") +

labs(

x = "Bootstrap P value",

y = "True P value",

title = "ZIP"

) +

geom_abline(intercept = 0, slope = 1) +

theme_classic()+

annotate(

geom = "text",

x = min(result$ZIP_synergy_p_value),

y = max(result$true_pv) - 0.1,

label = paste0(

"r = ", signif(c$estimate, 2),

"\nP = ", signif(c$p.value, 2)

),

hjust = 0

) +

theme(

panel.background = ggplot2::element_rect(

fill = "white",

colour = "white",

size = 2,

linetype = "solid"

),

panel.grid.major = ggplot2::element_line(

size = 0.5,

linetype = 'solid',

colour = "#DFDFDF"

),

panel.grid.minor = ggplot2::element_line(

size = 0.25,

linetype = 'solid',

colour = "#DFDFDF"

),

plot.title = ggplot2::element_text(

size = 13.5,

face = "bold",

hjust = 0.5

),

axis.text = ggplot2::element_text(

size = 10,

color = "black"

),

axis.title = ggplot2::element_text(

size = 10,

face = "italic"

),

\# axis.ticks = element_blank(),

axis.line.y.left = ggplot2::element_line(color = "black"),

axis.line.x.bottom = ggplot2::element_line(color = "black")

)

ggsave(

paste0(output_dir, "Scatter_plot_for_p_values.png"),

p, width = 3.5, height = 3

)

\# Loewe score -------------------------------------------------------------

\# Clean the environment

common_variables <- c(

"p_a", "p_b", "n", "dose", "k", "dose_comb",

"v", "s", "n_bootstrap", "N", "f_ll", "translate_p_value"

)

rm(list = setdiff(ls(), common_variables))

\# Output directory

output_dir <- paste0("../data/Loewe_synergyfinder_", n_bootstrap, "/")

if (!dir.exists(output_dir)){

dir.create(output_dir, recursive = TRUE)

}

\## Define functions for Loewe model ---------------------------------------

#' Solve the Expected Dose of Drug to Achieve Given Effect from LL.4 Model

#'

#' This function will solve the fitted four-parameter log-logistic dose-response

#' model and output the dose of drug at which it could achieve the \% inhibition

#' to cell growth.

#'

#' @param y The expected effect (\% inhibition) of the drug to cell line

#' @param drug_par The parameters for fitted dose-response model.

#'

#' @return A numeric value. It indicates the expected dose of drug.

#'

#' @author

#'	\itemize{

#'	}

#'

SolveExpDoesLL4 <- function(y, drug_par){

res <- drug_par[4] * (((y - drug_par[2]) / (drug_par[1] - y)) ^ (-1/drug_par[3]))

\# if NAN it means the response cannot be achieved by the drug, the response

\# can be either too high or too low for the drug to achieve

if (is.nan(res) == TRUE){

res <- ifelse(y > max(drug_par[2], drug_par[1]), Inf, 0)

}

return(res)

}

#' Solve the Loewe Additive Effect for Concentration Combinations Isobologram

#'

#' @param concs A numeric vector. It contains the concentrations of tested

#'	drugs.

#' @param drug_par A numeric vector. The parameters in fitted dose response

#'	curve.

#' @param drug_type The type of model used to fit dose response curve.

#' @param nsteps The total steps to calculate concentration combinations

#'	approaching to the true Loewe effect.

#'

#' @return A list contains 3 items:

#'	\itemize{

#'	\item y_loewe the predicted Loewe additive effect which closes to .

#'	\item x_select the expected concentrations for each drug to achieve

#'	y_loewe.

#'	\item distance the smallest distance

#'	}

#'

#' @author

#'	\itemize{

#' }

#'

SolveLoewe <- function(concs, drug_par, nsteps = 100){

\# x is the concentration vector

x <- as.matrix(concs)

dist <- rep(0, nsteps)

x_test <- mat.or.vec(2, nsteps)

y_test <- seq(0, 1, length.out = nsteps) # test nsteps responses

\# Calculate expected dose at each drug

for(i in 1:2){

tmp <- lapply(

y_test,

function(y) {

SolveExpDoesLL4(

y,

drug_par = drug_par[[i]]

)

}

)

x_test[i,] <- unlist(tmp)

}

\# Calculate distance between point x to the expected dose plane

for(j in 1:nsteps){

\# note the sign of -1

\# dist[j] = .Distance(w = c(1 / x_test[, j]), b = -1, point = x)

dist[j] = abs(sum(x/x_test[, j]) - 1)

}

index <- which(dist == min(dist, na.rm = T))

index <- min(index)

\# output the y_loewe corresponding to the minimal distance

res <- list(

y_loewe = y_test[index],

x_select = x_test[,index],

distance = min(dist, na.rm = T)

)

return(res)

}

\# model for inhibition (hill coefficient positive)

f_loewe <- function(x, p_a, p_b){

y <- SolveLoewe(

concs = c(x[1], x[2]),

drug_par = list(p_a, p_b)

)

return(y$y_loewe)

}

\## Expected response values -----------------------------------------------

\# The mono-therapy expected values (log-logistic dose-response curve)

m_a <- rep(f_ll(dose, p_a), each = n)

m_b <- rep(f_ll(dose, p_b), each = n)

\# The combo-therapy expected values (H0: Loewe model)

m_ab <- rep(apply(dose_comb, 1, f_loewe, p_a, p_b), each = n)

\## Distribution under null hypothesis -------------------------------------

\# Simulating the distribution of the test statistic under the null hypothesis

t_sample <- rep(0, N)

for (i in seq_len(N)) {

\# simulate mono-therapy data

y_a <- rnorm(n = length(m_a), mean = m_a, sd = s)

y_b <- rnorm(n = length(m_b), mean = m_b, sd = s)

\# simulate combo-therapy data

y_ab <- rnorm(n = length(m_ab), mean = m_ab, sd = s)

\# compute the sample means for each dose/combination

obs_m_a <- c(by(y_a, rep(seq_len(k), each = n), mean))

obs_m_b <- c(by(y_b, rep(seq_len(k), each = n), mean))

obs_m_ab <- c(by(y_ab, rep(seq_len(v), each = n), mean))

\# cross the previous sample means to compute the loewe value

response <- data.frame(

conc1 = c(0, dose, rep(0, k), dose_comb[, "Var1"]),

conc2 = c(0, rep(0, k), dose, dose_comb[, "Var2"]),

response = c(0, obs_m_a, obs_m_b, obs_m_ab) * 100,

stringsAsFactors = FALSE

)

synsco <- synergyfinder::Loewe(response)

synsco <- mean(synsco$Loewe_synergy[(2*k+2):nrow(synsco)])/100

\# average of the synergy score over the whole matrix

t_sample[i] <- synsco

}

\## Bootstrap --------------------------------------------------------------

\# Bootstrap function for Loewe model

bootstrap <- function(i, m_a, m_b, m_ab, s, N, dose_comb, k, t_sample){

set.seed(i)

\# generate a new sample for the bootstrap study

y_a <- rnorm(n = length(m_a), mean = m_a, sd = s)

y_b <- rnorm(n = length(m_b), mean = m_b, sd = s)

y_ab <- rnorm(n = length(m_ab), mean = m_ab, sd = s)

\# compute the observed statistics

obs_m_a <- c(by(y_a, rep(seq_len(k), each = n), mean))

obs_m_b <- c(by(y_b, rep(seq_len(k), each = n), mean))

obs_m_ab <- c(by(y_ab, rep(seq_len(v), each = n), mean))

response <- data.frame(

conc1 = c(0, dose, rep(0, k), dose_comb[, "Var1"]),

conc2 = c(0, rep(0, k), dose, dose_comb[, "Var2"]),

response = c(0, obs_m_a, obs_m_b, obs_m_ab) * 100,

stringsAsFactors = FALSE

)

obs_Loewe <- synergyfinder::Loewe(response)

t_obs <- mean(obs_Loewe$Loewe_synergy[(2*k+2):nrow(obs_Loewe)])/100

\# true p-value

true_pv <- mean((t_sample <= -abs(t_obs)) | (t_sample > abs(t_obs)))

\# do bootstrap...

n_well <- (1+2*k+nrow(dose_comb)) * n

data_table <- data.frame(

block_id = rep(1, times = n_well),

drug1 = rep("drug_a", times = n_well),

drug2 = rep("drug_b", times = n_well),

conc1 = rep(c(0, dose, rep(0, k), dose_comb[, "Var1"]), each = n),

conc2 = rep(c(0, rep(0, k), dose, dose_comb[, "Var2"]), each = n),

ConcUnit = rep("uM", times = n_well),

response = c(rep(0, times = n), y_a, y_b, y_ab) * 100,

stringsAsFactors = FALSE

)

data <- synergyfinder::ReshapeData(data_table, data_type = "inhibition")

res <- synergyfinder::CalculateSynergy(

data,

iteration = N,

seed = i,

method = "Loewe"

)

boot_pv <- res$drug_pairs

boot_pv$iteration = i

boot_pv$true_pv <- true_pv

\# plot(density(t_sample), xlim = c(-0.05, 0.05))

\# plot(density(t_boot), xlim = c(-0.05, 0.05))

return(

p_values = boot_pv

)

}

\# Run bootstrap

result <- NULL

for (i in 1:n_bootstrap) {

tmp <- bootstrap(

job_id, m_a, m_b, m_ab, s,

N = n_bootstrap, dose_comb, k, t_sample

)

result <- rbind.data.frame(result, tmp)

}

if (!is.numeric(result$Loewe_synergy_p_value)){

result$Loewe_synergy_p_value <- sapply (

result$Loewe_synergy_p_value,

translate_p_value

)

}

write.csv(result, paste0(output_dir, "P_value.csv"), row.names = FALSE)

\## Visualization ----------------------------------------------------------

c <- cor.test(result$Loewe_synergy_p_value, result$true_pv)

p <- ggplot(result, aes(x = Loewe_synergy_p_value, y = true_pv)) +

geom_point(color = "#2D72AD") +

labs(

x = "Bootstrap P value",

y = "True P value",

title = "Loewe"

) +

geom_abline(intercept = 0, slope = 1) +

theme_classic()+

annotate(

geom = "text",

x = min(result$Loewe_synergy_p_value),

y = max(result$true_pv) - 0.1,

label = paste0("r = ", signif(c$estimate, 2), "\nP = ", signif(c$p.value, 2)),

hjust = 0

) +

theme(

panel.background = ggplot2::element_rect(

fill = "white",

colour = "white",

size = 2,

linetype = "solid"

),

panel.grid.major = ggplot2::element_line(

size = 0.5,

linetype = 'solid',

colour = "#DFDFDF"

),

panel.grid.minor = ggplot2::element_line(

size = 0.25,

linetype = 'solid',

colour = "#DFDFDF"

),

plot.title = ggplot2::element_text(

size = 13.5,

face = "bold",

hjust = 0.5

),

axis.text = ggplot2::element_text(

size = 10,

color = "black"

),

axis.title = ggplot2::element_text(

size = 10,

face = "italic"

),

\# axis.ticks = element_blank(),

axis.line.y.left = ggplot2::element_line(color = "black"),

axis.line.x.bottom = ggplot2::element_line(color = "black")

)

ggsave(paste0(output_dir, "Scatter_plot_for_p_values.png"), p,

width = 3.5, height = 3)

**Session information for testing**

R version 4.1.1 (2021-08-10)

Platform: x86_64-apple-darwin17.0 (64-bit)

Running under: macOS Catalina 10.15.7

Matrix products: default

BLAS: /System/Library/Frameworks/Accelerate.framework/Versions/A/Frameworks/vecLib.framework/Versions/A/libBLAS.dylib

LAPACK: /Library/Frameworks/R.framework/Versions/4.1/Resources/lib/libRlapack.dylib

locale:

[1] en_US.UTF-8/en_US.UTF-8/en_US.UTF-8/C/en_US.UTF-8/en_US.UTF-8

attached base packages:

[1] stats	graphics grDevices utils	datasets methods	base

other attached packages:

[1] ggplot2_3.3.5	synergyfinder_3.0.15

loaded via a namespace (and not attached):

[1] gtools_3.9.2	zoo_1.8-9	tidyselect_1.1.1 purrr_0.3.4	splines_4.1.1	haven_2.4.3	lattice_0.20-45	carData_3.0-4

[9] colorspace_2.0-2 vctrs_0.3.8	generics_0.1.0	utf8_1.2.2	survival_3.2-13	rlang_0.4.11	pillar_1.6.2	withr_2.4.2

[17] foreign_0.8-81	glue_1.4.2	DBI_1.1.1	readxl_1.3.1	multcomp_1.4-17	lifecycle_1.0.1	munsell_0.5.0	gtable_0.3.0

[25] cellranger_1.1.0 zip_2.2.0	mvtnorm_1.1-2	codetools_0.2-18 rio_0.5.27	forcats_0.5.1	curl_4.3.2	fansi_0.5.0

[33] TH.data_1.0-10	Rcpp_1.0.7	scales_1.1.1	plotrix_3.8-2	abind_1.4-5	hms_1.1.0	stringi_1.7.4	openxlsx_4.2.4

[41] dplyr_1.0.7	grid_4.1.1	tools_4.1.1	sandwich_3.0-1	magrittr_2.0.1	tibble_3.1.5	crayon_1.4.1	car_3.0-11

[49] drc_3.0-1	pkgconfig_2.0.3	ellipsis_0.3.2	MASS_7.3-54	Matrix_1.3-4	data.table_1.14.0 assertthat_0.2.1 R6_2.5.1

[57] compiler_4.1.1
